# Locations of consecutive G•C base-pairs direct genomic nucleosome positioning

**DOI:** 10.1101/2025.09.01.673439

**Authors:** Hiroaki Kato, Tomohiro Fuse, Shoko Sato, Yohei Kurihara, Wataru Kagawa, Takeshi Urano, Yasuyuki Ohkawa, Hitoshi Kurumizaka, Mitsuhiro Shimizu

**Affiliations:** Department of Biochemistry, Shimane University School of Medicine, Izumo, Shimane, 693-8501, Japan; Department of Chemistry, Graduate School of Science and Engineering, Meisei University, 2-1-1 Hodokubo, Hino, Tokyo 191-8506, Japan; Laboratory of Chromatin Structure and Function, Institute for Quantitative Biosciences, The University of Tokyo, 1-1-1 Yayoi, Bunkyo-ku, Tokyo 113-0032, Japan; Division of Transcriptomics, Medical Institute of Bioregulation, Kyushu University, 3-1-1 Maidashi, Higashi, Fukuoka 812-0054, Japan

## Abstract

The nucleosome is a structural and functional subunit of chromatin, and its positioning and dynamics in the eukaryotic genomes serves as a key platform for gene regulation. Here, we determine the positions of fully wrapped nucleosomes across the yeast genome by chemical mapping through the histone H2A-A122C residue, which cleaves near the DNA entry/exit sites. This approach reveals the most refined sequence-dependent profile of nucleosomes reported to date. Comparison of H2A-A122C with the H3-Q85C and H4-S47C methods clearly shows a sequence preference for the chemical cleavage site, and mapping results suggest that the nucleosome locations are dynamic *in vivo*. More importantly, we find that the depletion of CC and GG dinucleotides at nucleotide positions -11 to -9 and +9 to +11 bp, respectively, from the nucleosome dyad position (0) is inextricably associated with the enrichment of AA/AT/TA/TT dinucleotides in both yeast and mouse genomes. Introducing consecutive C•G base pairs to the corresponding sites in the 601 sequence shifts the nucleosomes to less frequent positions without altering thermal stability, implying a structural constraint imposed by DNA sequence. Thus, CC and GG dinucleotides in the major groove blocks at superhelix locations (SHL) -1.0 and +1.0, respectively, destabilize histone-DNA interactions, serving as intrinsic determinants of nucleosome positioning in eukaryotic genomes.

## Introduction

The nucleosome is a structural and functional unit in eukaryotic genomes, in which 145-147 bp of DNA are wrapped in ∼1.7 left-handed superhelical turns around the histone octamer (two molecules each of histones H2A, H2B, H3 and H4) (Luger et al. 1997; White et al. 2001; reviewed in McGinty and Tan 2015; Takizawa and Kurumizaka 2022). Understanding the determinants of nucleosome positioning *in vivo* is important for elucidating chromatin function and epigenetic regulation (reviewed in Struhl and Segal 2013; Lai and Pugh 2017; Chereji and Clark 2018; Brahma and Henikoff 2020; Singh and Mueller-Planitz 2021; Baldi et al. 2020). Many methods have been developed to determine nucleosome locations, and site-directed chemical mapping *via* histones provides high resolution to genomic coordinates (reviewed in Voong et al. 2017). Serine 47 of histone H4 was substituted with cysteine (H4-S47C) to link with a copper-chelating reagent, and hydroxy radicals were locally generated by the addition of hydrogen peroxide to cleave the DNA backbones around the dyad of the nucleosome (Brogaard et al. 2012a). The cleaved H4-S47C fragments were then subjected to massive parallel sequencing. This method was successfully applied to the genome-wide mapping of nucleosomes in *S. cerevisiae*, *S. pombe* and mouse embryonic stem cells (Brogaard et al. 2012a; Moyle-Heyrman et al. 2013; Henikoff et al. 2014; Voong et al. 2016; Fuse et al. 2017; Chereji et al. 2017; Fuse et al. 2019; Katsumata et al. 2021). However, the definition of the nucleosome center position (dyad) was not straightforward, due to the complexity of the cleavage patterns around the dyad, and a Bayesian deconvolution algorithm was developed to calculate the nucleosome center position score (Brogaard et al. 2012a). Importantly, the H4-S47C cleavage may have a minor sequence bias (Brogaard et al. 2012a), and our previous studies also showed that H4-S47C preferentially cleaves at (AAT)_12_ and (ACT)_12_ sequences at the center of nucleosomes (Katsumata et al. 2021). Subsequently, the histone H3-Q85C chemical mapping was developed (Chereji et al. 2018). The nucleosomal dyad was more simply deduced from the center of the 51 bp fragment obtained from the H3-Q85C-cleavages at SHL (SuperHelix Location) ±2.5 from the dyad of the nucleosome. Thus, the H3-Q85C method provided precise mapping of both nucleosomes and linkers in *S. cerevisiae*. Since H4-S47C and H3-Q85C cleave near SHL ±0.5 and ±2.5, respectively, these methods might detect not only fully wrapped nucleosomes but also unwrapped nucleosomes. Further clarification of the role of DNA sequences in nucleosome positioning requires the development of new high-resolution mapping methods for fully wrapped nucleosomes, preferably with unbiased intranucleosomal sequences.

Herein, we describe a novel chemical mapping method using the H2A-A122C residue, which cleaved the entry/exit sites of a single nucleosome, resulting in unbiased intranucleosomal sequences. Comparisons of H2A-A122C mapping data with H4-S47C, H3-Q85C, and MNase-seq data revealed the sequence dependency of the nucleosome dynamics *in vivo* at a higher resolution. Furthermore, the H2A-A122C data demonstrated the enrichment of the WW (AA/AT/TA/TT) dinucleotide distribution in the regions of -10 to -8 and +8 to +10 relative to the dyad, and the concomitant scarcities of CC and GG dinucleotides at -11 to -9 and +9 to +11, respectively. The introduction of CC and/or GG dinucleotides into the 601-nucleosome showed that consecutive G•C base pairs at these critical sites destabilize the nucleosome position. A unifying feature for DNA sequence-dependent nucleosome formation *in vivo* is proposed.

## Results

### Genome-wide chemical mapping of nucleosomes with H2A-A122C

The crystal structures, all-atom MD simulations, and cross-linking studies of nucleosomes have shown that the C-terminal tail of histone H2A protrudes from the nucleosome core particles and interacts with both the nucleosomal and linker DNA regions (Luger et al. 1997; Lee and Hayes 1998; Ghoneim et al. 2021; Li and Kono 2016; Armeev et al. 2021). We introduced the mutation of Ala122 (amino acid numbering except for the first methionine at the start of translation) to cysteine in histone H2A (H2A-A122C) of *S. cerevisiae* to apply site-directed chemical mapping **(Fig. 1A and Supplemental Figs. S1 and S2**). Analysis of the cleavage sites by high-throughput sequencing revealed that the nucleotides located at ±70 nucleotides from the dyad of the nucleosomal DNA were mainly cleaved, releasing ∼139 bp fragments (**Figs. 1B and 1C**). This analysis confirmed that H3-Q85C and H4-S47C cleaved at ±2 and ±5 and ±26 and ±29 nucleotides from the dyad, respectively, consistent with previous studies (Brogaard et al. 2012a; Chereji et al. 2018; Fuse et al. 2017) (**Supplemental Fig. S2A**). Cleavage by H2A-A122C at ±73 nucleotides was also detected for DNA fragments that extended outward, and peaked at 157 and 167 bp lengths (**Fig. 1B**). Considering the gap width between the inner and outer fragments, the major nucleosome repeat length estimated by this method (**Supplemental Fig. S2B**) was in good agreement with previous studies (Cui and Zhurkin 2010; Brogaard et al. 2012a; Chereji et al. 2018).

**Fig. 1.**
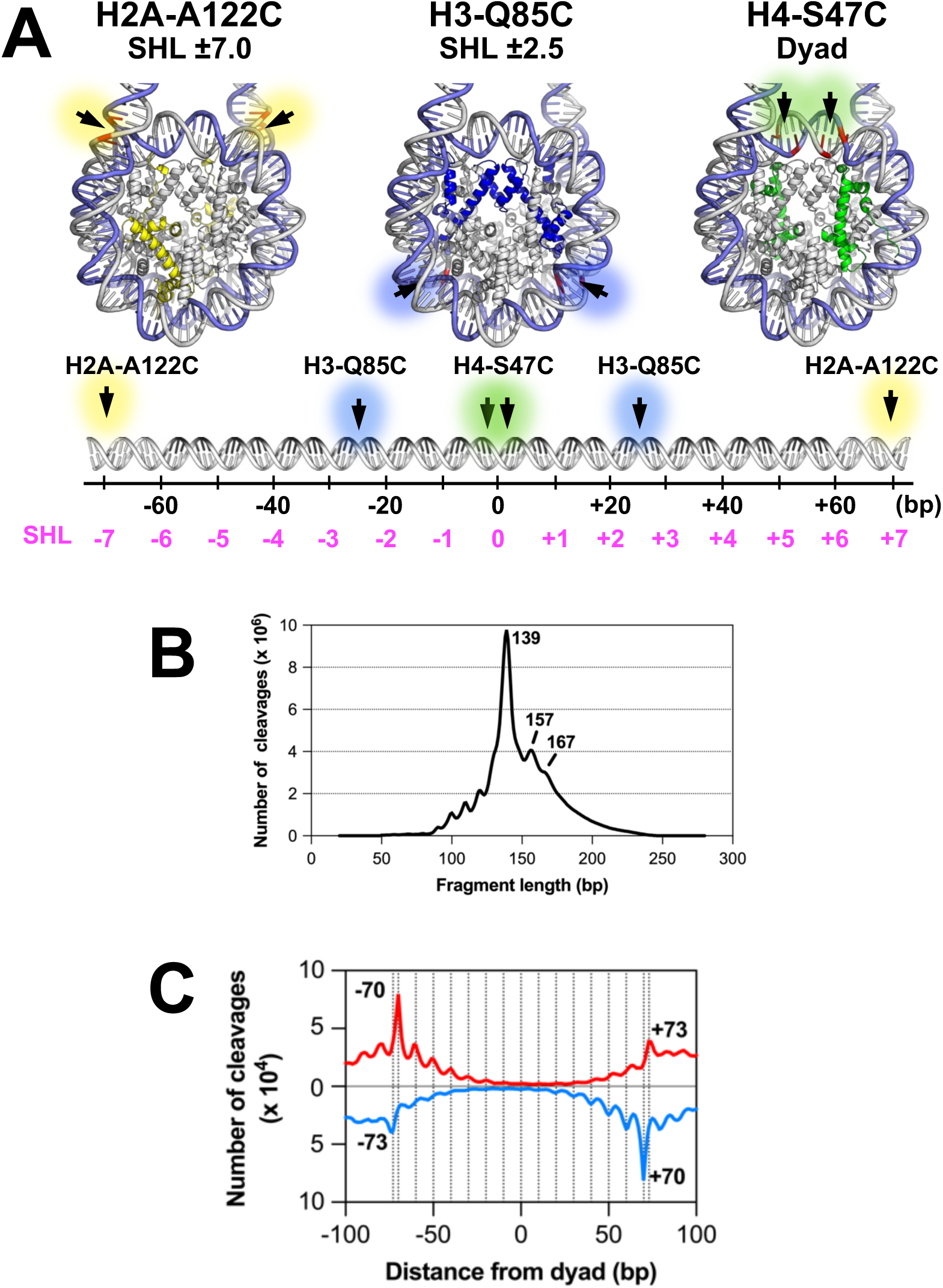
Site-specific chemical cleavages of nucleosomal DNA. **(A)** Schematic representation of chemical cleavage sites in nucleosomal DNA *via* histone H2A-A122C, H3-Q85C or H4-S47C residues linked to the copper-chelating label, N-(1,10-phenanthroline-5-yl) iodoacetamide. Crystal structure of the yeast nucleosome (PDB: 1ID3)(White et al., 2001), with histones H2A, H3 and H4 colored yellow, blue and green, respectively. Each cleavage site is represented on the DNA duplex. SHL: superhelix location. **(B)** Lengths of DNA fragments produced by the H2A-A122C chemical cleavages. These fragments were subjected to next-generation sequencing. **(C)** Occurrences of H2A-A122C cleaved sites in 66,434 representative nucleosomes in the yeast genome. DNA fragments from the representative nucleosomes were aligned with each dyad position as zero.

To confirm the reliability of the chromatin structure of the H2A-A122C strain, we performed MNase-seq analyses of the H2A-A122C and wild-type strains (**Supplemental Figs. S1 and S3**). The results showed that our samples did not undergo extensive digestion, with most mapped fragments over 147 bp in length and peaking at 153, 162 and 170 bp (**Supplemental Fig. S3A)**. The MNase-cleaved profiles for the representative 67,510 nucleosomes in the yeast genome (**Supplemental Fig. S3B**), and for the -1 to +6 nucleosomes in 5,542 genes (**Supplemental Fig. S3C**) were in good agreement between the H2A-A122C and wild type strains. Accordingly, the introduction of the H2A-A122C mutation has little or no effect on the formation and positioning of nucleosomes *in vivo*.

To determine the nucleosome positions in the yeast genome by H2A-A122C chemical mapping, we counted the numbers of 139±5 bp mapped fragments at their midpoint on the genomic coordinates as the “dyad score”, as conducted in the H3-Q85C method (Chereji et al. 2018). For the H2A-A122C and H3-Q85C methods, the genomic coordinates of the highest dyad score within a given ±107 bp region were chosen as the position of the dyad base of a representative nucleosome. The allowance of nucleosomal overlap (40 bp at most for the neighboring nucleosome wrapped by 147 bp DNA) was previously applied for the H4-S47C-based nucleosome calling (Brogaard et al. 2012a). The dyad-to-dyad distance between representative nucleosomes peaked at 162±10 bp for the three methods (**Supplemental Fig. S2B**). We performed H4-S47C-based mapping of independent biological samples of the strain with the genetic background of FY24 (Fuse et al. 2017) to confirm the reliability of our methodology. The positions of representative nucleosomes identified by the H4-S47C method in this study matched well with those determined previously (Brogaard et al. 2012a) (**Supplemental Fig. S4**). As for the H2A-A122C method, independent experiments confirmed that the nucleosome calling with this method was highly reproducible (**Supplemental Fig. S4A, Upper panel**). As a result, we obtained a set of representative nucleosomes for evaluating the three kinds of chemical methods. Since H3-Q85C and H4-S47C cleaved around SHLs ±2.5 and ±0.5, respectively, it is not clear whether the DNA binds histone octamers towards the entry/exit sites of the nucleosomes. Thus, it is worth noting that the H2A-A122C method can detect fully wrapped nucleosomes because it cleaves near SHL ±7.0.

### Comparison of H2A-A122C with the H3-Q85C and H4-S47C mappings

To clarify the sequence-dependence of nucleosome formation, we examined the distribution of 16 dinucleotides within representative nucleosomes determined by H2A-A122C, and compared it with those of H3-Q85C and H4-S47C (**Figs. 2A and B and Supplemental Fig. S5**). This comparison clearly revealed that H4-S47C-based nucleosomes tend to have extraordinarily high proportions of AC/GT, AG/CT and TA around the dyad (red bars in lower panel of **Fig. 2A**), consistent with a previous report (Brogaard et al. 2012a). As for those identified by the H3-Q85C method, biases are also detected around SHL ±2.5 (**Fig. 2A**, gray shadow in **Fig. 2B)**. In the case of H2A-A122C-based nucleosomes, AA-TT dinucleotides are strongly disfavored at the terminal regions of nucleosomal DNA where cleavage occurs (SHL ±7) (**Fig. 2A, gray shadow in Fig. 2B and Supplemental Fig. S5**). Apart from the terminal bias at SHL ±7, the dinucleotide frequency of the H2A-A122C-based nucleosomes did not appear to be biased in the inner region (SHL –6 to +6). Based on the 16 dinucleotide-frequency heat maps (**Fig. 2A**), the WW/SS (W=A or T, S=G or C), RR/YY (R=A or G, Y=C or T) and RY/YR frequencies were plotted (**Fig. 2B and Supplemental Fig. S5**). The RR/YY and YR/RY plots clearly showed that each chemical method has preferences for particular dinucleotides around the cleavage sites (gray shadows in **Supplemental Figs. S5B and S5C**).

**Fig. 2.**
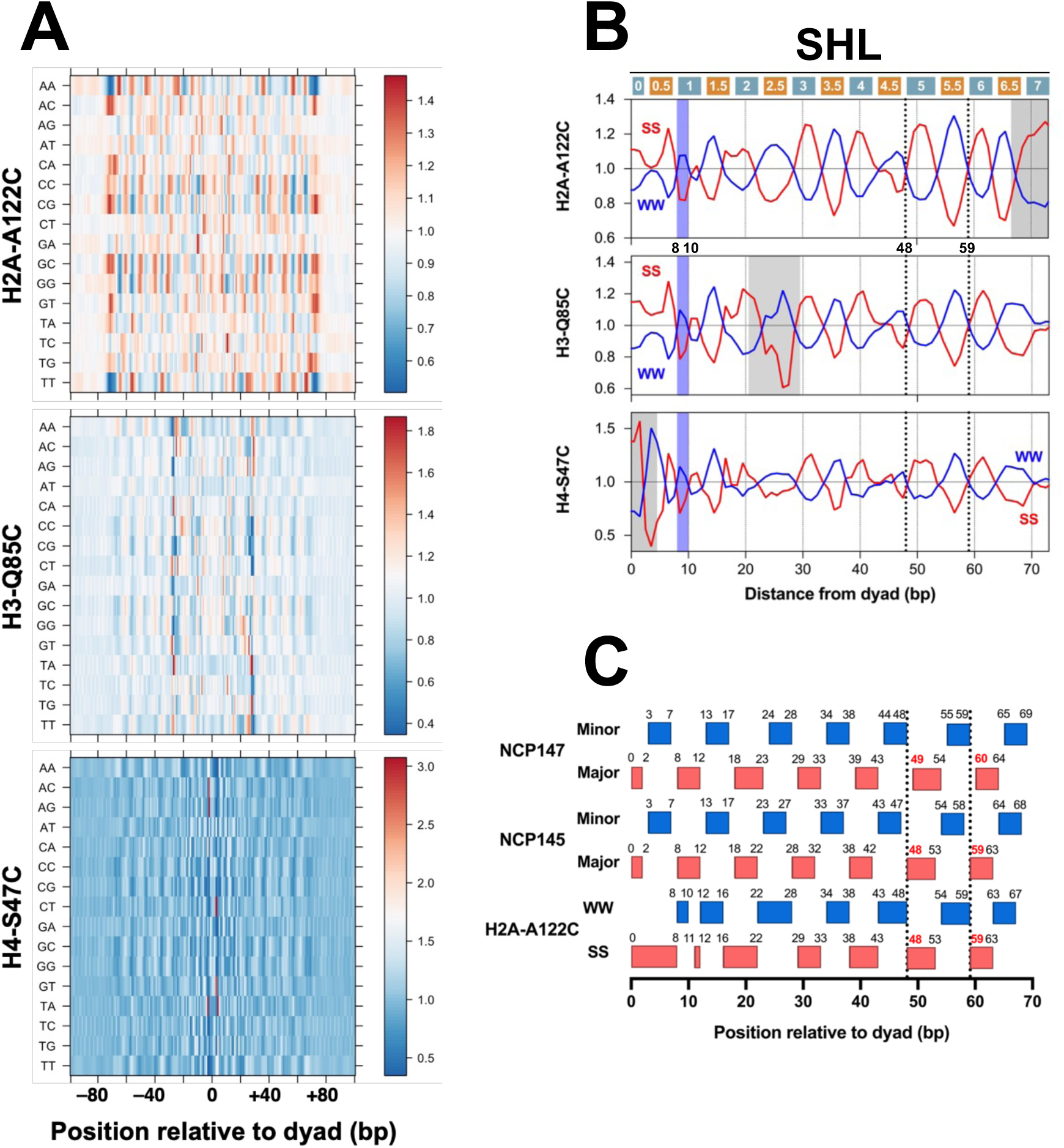
Genome-wide analysis of DNA sequence-dependence of representative nucleosomes in yeast. DNA sequences from the nucleosomes were aligned with each dyad position as 0. **(A)** Heat map analysis of the occurrence of 16 dinucleotides within the representative nucleosomes (H2A-122C: 66,434, H3-Q85C: 60,559, H4-S47C: 67,510) in the yeast genome, whose positions were determined by H2A-A122C (this study), as well as the previously reported H3-Q85C (Chereji et al. 2018) and H4-S47C (Brogaard et al. 2012) chemical mappings. **(B)** Frequency of WW (AA, TT, AT, TA) and SS (GG, CC, GC, CG) dinucleotides, shown in blue and red, respectively, in the representative nucleosomes in yeast as determined by H2A-A122C, H3-Q85C or H4-S47C chemical mapping. One side of the nucleosomes (0 to +75) is shown since the profile is symmetrical with regard to the dyad. The whole regions of these profiles are shown in Figure S5A. The light blue shaded regions indicate the WW sites at around SHL +1.0. The gray shaded regions indicate sequence biases at each chemical cleaved site. The dotted lines correspond to the dotted lines in Fig. 2C**. (C)** Comparison of the WW/SS frequency distributions in representative nucleosomes in yeast with the positions of major and minor groove blocks in the crystal structures of nucleosomes (PDBs: 1KX5, 2NZD(Ong et al. 2007, Richmond and Darvey 2003)) containing 147 bp or 145 bp DNA (denoted NCP147 and NCP145, respectively).

The H2A-A122C, H3-Q85C and H4-S47C methods determined the positions of representative nucleosomes as 66,434, 60,559 and 67,510, respectively. We compared the positions of the representative nucleosomes identified by the three types of chemical cleavages (**Supplemental Fig. S6A**). The coincidence at ±1 bp resolution of the identified dyad positions between any two chemical methods was about 10-15%, and the population peaks of the other fractions showed symmetry at about 10 bp periodicities. These results support the idea that the translational position of nucleosomes changes dynamically throughout the yeast genome while maintaining the DNA surface contacts with histones. This is consistent with previous reports (Buttinelli et al. 1993; Kim et al. 2006; Shen et al. 2001) that multiple translational positionings by a unique helical repeat were observed in the *5S rRNA*, *CUP1* and *HIS3* genes in *S. cerevisiae*. We also calculated the intersections of representative dyad positions determined by the three methods and drew their Venn diagrams (**Supplemental Fig. S6B**). The subset of populations of nucleosome positions determined by each method had approximately the same range. Thus, it is considered that while the positions of the nucleosome change dynamically *in vivo*, each chemical method cleaves the DNA when it encounters the preferred sequence among the overlapping and redundant positions of nucleosomes. The preference may be determined by the local affinities of the histone and the copper chelate label, N-(1,10-phenanthroline-5-yl)iodoacetamide, for the cleaved sites.

### Sequence-dependence within fully wrapped nucleosomes *in vivo*

The WW/SS profile of the H2A-A122C-based nucleosomes is similar to those of the H3-Q85C-based and H4-S47C-based nucleosomes, except for the regions of their cleaved sites (gray shadows in **Fig. 2B and Supplemental Fig. S5A**). Thus, the H2A-A122C method provides the most unbiased view of dinucleotide frequency in the inner region of nucleosomal DNA, since it cuts the entry/exit sites. These results are consistent with previous studies showing that the SS and WW dinucleotides peaked in the major groove and minor groove blocks, respectively, in the nucleosome.

In addition, the analysis of the H2A-A122C-based nucleosome revealed ambiguities in the details of the WW/SS profile. Contrary to the general trend, it should be noted that there is a WW peak at around SHL ±1.0, as shown by light blue shadings in Figs. 2B and S5A. The WW sites at the region of -10 to -8 bp and +8 to +10 bp from the dyad would be important to contact the alpha N helix of histone H3 (Luger et al. 1997) and the N-terminal tail of H4 (Nikitina et al. 2025). The SS and WW peaks at SHL ±2 and ±2.5, respectively, are wider than those in other locations (**Fig. 2B and Supplemental Fig. S5A**). We also noticed that the SS sites at SHL ±4 apparently overlap with the WW sites at SHL ±4.5. This point is likely to be related with a recent *in vitro* reconstitution study showing that the interactions of the N-terminal tails of histones H2A with the SHL ±4.0 regions affect the sequence-dependent nucleosome formation (Nikitina et al. 2025). Interestingly, AA and TT dinucleotides exhibited direction-dependent distribution patterns at SHL ±2.5 and ±4.5 (**Fig. 2A and Supplemental Fig. S7D**), where AA/TT dinucleotides have long been known to exhibit heterogenic “camel-humped” patterns (Satchwell and Travers 1989). These position-specific features may reflect the locations of the H2B N-terminal tails that pass through the gyres of the DNA superhelix (Luger et al. 1997; Ohtomo et al. 2021). Additionally, as illustrated in the profile of the H2A-A122C of Fig. 2C, we note that the high SS frequency regions in the H2A-A122C-based nucleosomes at the major groove blocks 5 and 6 are located at 48-53 and 59-63 nucleotides from the dyad, respectively (dotted lines in **Fig. 2B and 2C**). Nucleosomes with this character are closer to NCP145, rather than NCP147 (Ong et al. 2007; Richmond and Davey 2003; Davey et al. 2002). Thus, there appear to be two occurrences of DNA stretching in the inner region of the wrapped DNA, and the majority is the 145 bp-NCP *in vivo*, which is consistent with a recent cryo-EM result that the NCP in solution is likely to include 145 bp of DNA (Bai and Zhou 2021).

### Lack of consecutive G•C pairs within SHL ±1.0

To examine the contribution of internal DNA sequences to nucleosome formation (DNA wrapping around histone octamers), we grouped the H2A-A122C-based representative nucleosomes by the dyad score, which reflects how often nucleosomes are detected at a particular genomic coordinate (**Supplemental Fig. S7A**). We also calculated the standard deviation for the relative positions of the surrounding nucleosomes from the representative ones, which indicates the fuzziness of the nucleosome position (Albert et al. 2007), and analyzed the correlation between the standard deviation and the dyad scores of representative nucleosomes (**Supplemental Fig. S7B**). The results showed that nucleosomes with higher dyad scores had lower SD values. This means that these nucleosomes tended to have lower positional redundancy, and consistently, the vertical stripe pattern of dinucleotides was observed more clearly in the group of nucleosomes with higher dyad scores (**Supplemental Fig. S7C**). These results also suggested that the biased region of H2A-A122C cleavage around the SHL ±7 terminal subsequence has little effect on the detection efficiency.

As expected from the WW/SS periodic pattern, in nucleosomes with higher dyad scores, CG, GC, CC and GG dinucleotides were prominent within the SS peaks in the major groove blocks in the WW/SS profile (**Fig. 3A and Supplemental Figs. S7C and S7D**). It is worth noting that CC dinucleotides were extremely rare in the -11 to -9 bp region of the SHL -1.0 major groove block, whereas GG dinucleotides were rare in the +9 to +11 bp region of the +1.0 major groove block (indicated by asterisks in **Fig. 3A).** This trend is independent of the nucleosome detection frequency, and inextricably linked to the fact that WW dinucleotides are preferred in these regions (light blue shading in **Fig. 2B**). We also analyzed previously reported chemical mapping data for H3-Q85C (Chereji et al. 2018) and H4-S47C (Brogaard et al. 2012a) to confirm our findings that CC dinucleotides are rarely found around positions -11 to -9 on yeast genome nucleosomes (middle and bottom panels in **Fig. 3A**). Since genome-wide mapping data by H4-S47C chemical cleavage are available for *S. pombe* (Moyle-Heyrman et al. 2013) and mouse embryonic stem cells (Voong et al. 2016), we further investigated the occurrence of CC dinucleotides in the nucleosomes of these species. CC dinucleotides were scarcely present in the -9 to -11 region in not only *S. cerevisiae* but also *S. pombe* and mouse, as shown in **Fig. 3B**. Thus, this position-dependent sequence preference in the nucleosome is conserved among different species.

**Fig. 3.**
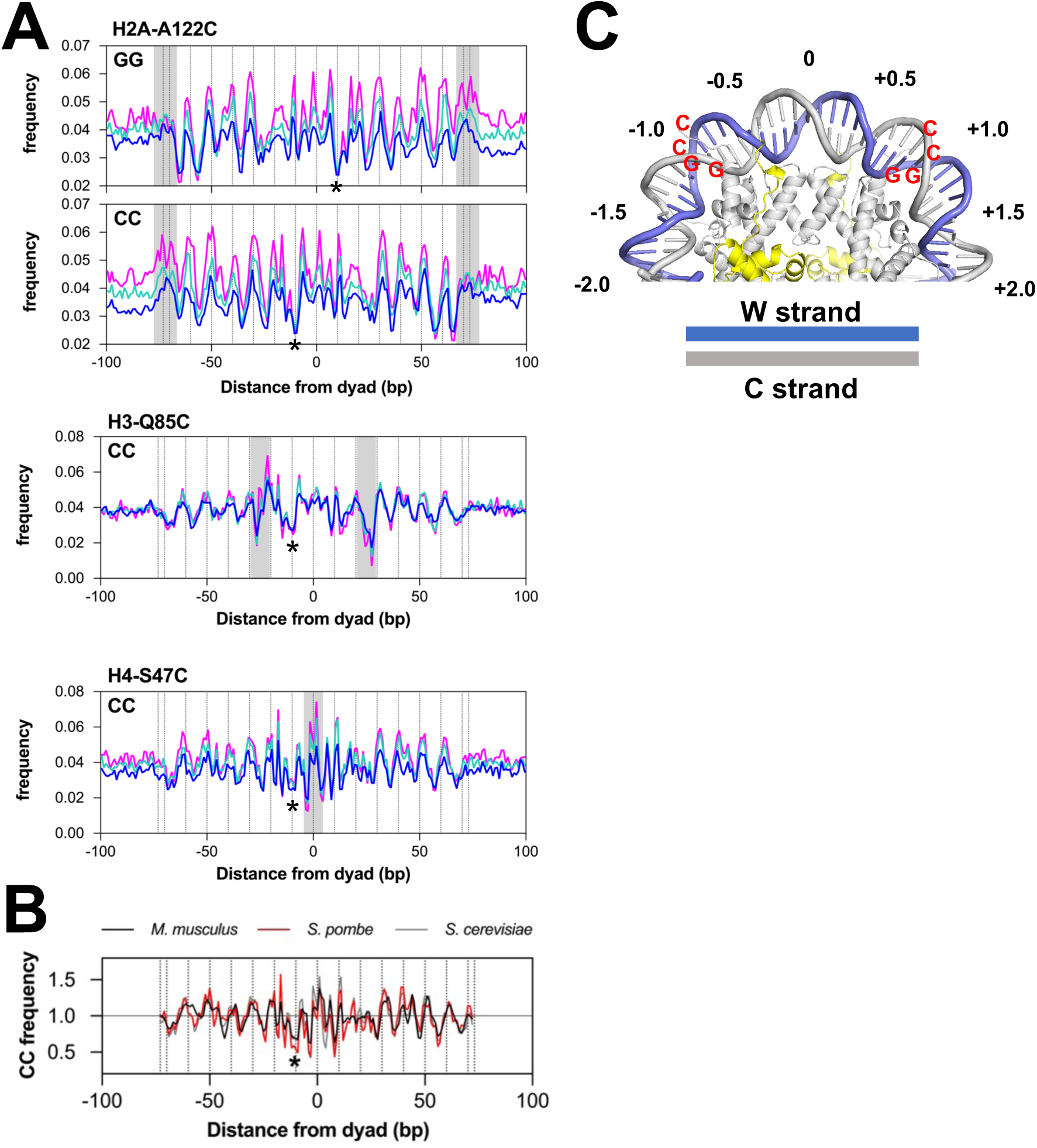
Occurrence of CC/GG dinucleotides within nucleosomes in the genomes. (A) Frequencies of CC and GG dinucleotides within representative nucleosomes in the yeast genome. Asterisks indicate sites where the frequencies of CC and GG dinucleotides are very rare. The representative nucleosomes determined by H2A-A122C chemical mapping were subdivided into nine groups of equal numbers, each according to the dyad score values (see Fig. S7A). The magenta, turquoise and blue lines indicate the results for groups 1, 5, and 9, respectively. Frequencies of CC dinucleotides from previously published results on chemical mapping of H3-Q85C (Chereji et al. 2017) or H4-S47C (Brogaard et al. 2012) are also shown. The magenta, turquoise and blue lines have the same meaning as the profiles obtained by H2A-A122C. **(B)** Frequency of CC dinucleotides analyzed from previously reported results on H4-S47C chemical mapping in *M. musculus* (black line), *S. pombe* (red line), and *S. cerevisiae* (gray line) (Brogaard et al. 2012, Moyle-Heyrman et al. 2013, Voong et al. 2016). **(C)** Locations of CC- and GG-rare sites in the nucleosomal DNA.

The rare CC and GG sites are symmetrical with respect to the dyad, which means that the C•G base pairs are oriented in the same direction in the DNA path on the histone octamer. As shown in **Fig. 3C**, the diagram of these rare sites on nucleosomal DNA shows that the guanine residues are on the proximal strand to the histone octamer, whereas the cytosine residues are on the distal strand. The presence of two consecutive guanine bases in the DNA strand proximal to the histone octamer may cause steric instability of the nucleosome.

### Effect of CC/GG substitution on nucleosome position *in vitro*

To examine the effect of consecutive C•G base pairs in the major groove block at SHL ±1.0 regions on nucleosome formation, we reconstituted nucleosomes *in vitro* using the 193 bp DNA fragment containing the 601 sequence and histone octamers (Sato et al. 2020). In this 193 bp fragment, the 601 sequence is located between nucleotides 27 and 167, and the crystal structure of the nucleosome suggests that the 96th cytosine residue on the top strand is the dyad base (Vasudevan et al. 2010) (**Supplemental Fig. S8A**). By mutating both the adenine at position -9 from the dyad to cytosine and the cytosine at position +10 to guanine, we designed a CC_mut1 fragment with CCC from -10 to 8 and GGG from +9 to +11 (**Supplemental Fig. S8B**). We also constructed each single mutation, CC_mut1_L with CCC mutations from -10 to -8 and CC_mut1_R with GGG from +9 to +11 **(Fig. 4A**). We reconstituted nucleosomes with histone octamers and 193 bp fragments of either 601, CC_mut1, CC_mut1_L or CC_mut1_R by the salt-dialysis method (**Supplemental Fig. S9A**). The nucleosomes were purified by preparative PAGE (polyacrylamide gel electrophoresis), and their integrities were checked by native PAGE and SDS-PAGE (**Supplemental Figs. S9B and S9C**).

**Fig. 4.**
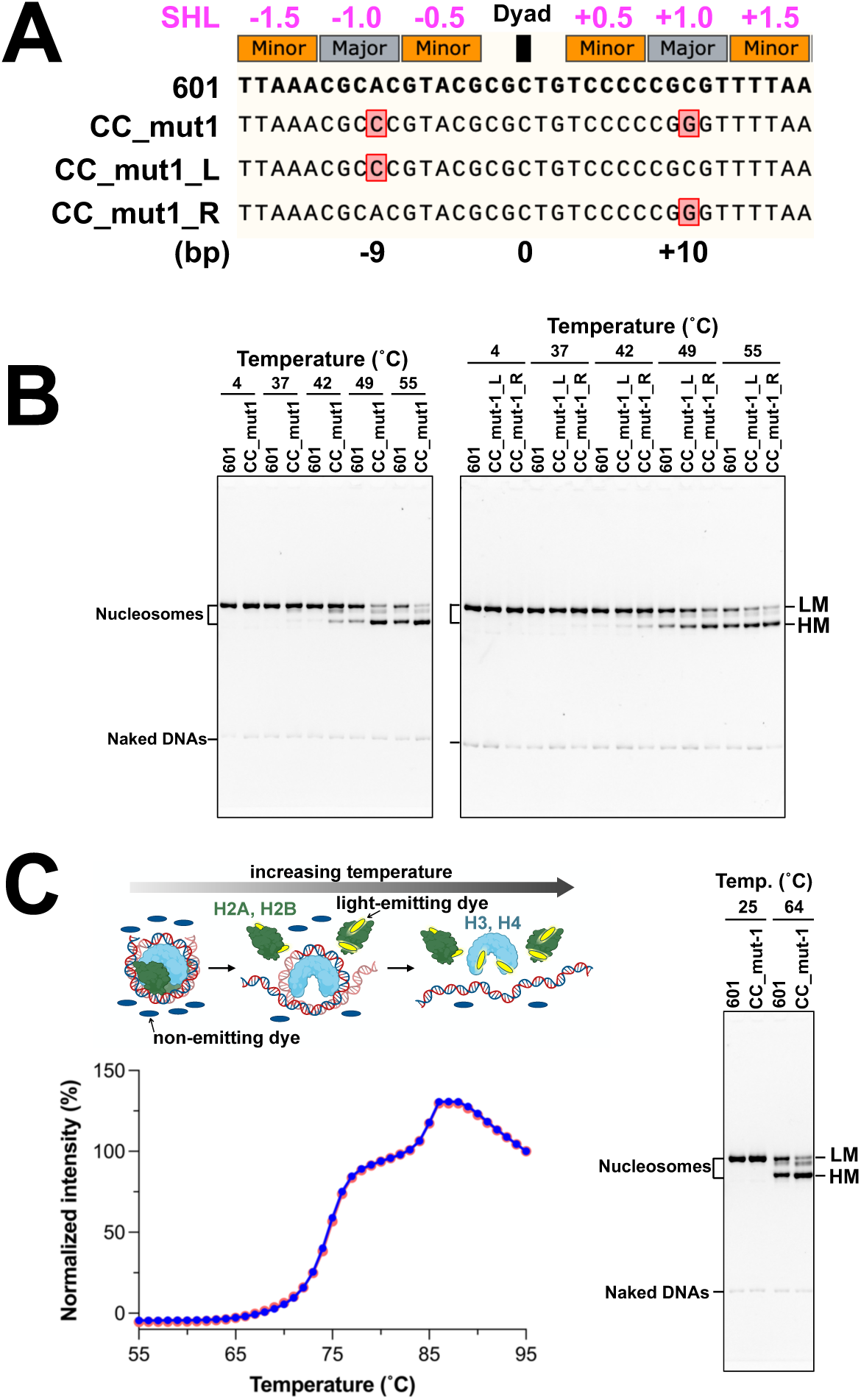
Effect of CC/GG mutations on nucleosome positioning in the 601-nucleosome reconstituted *in vitro*. **(A)** Design of DNA mutation sites based on the crystal structure of the 601-nucleosome (PDB: 3LZ0)(Vasudevan et al. 2010). In CC_mut1, adenine at -9 and cytosine at +10 from the dyad were replaced by cytosine and guanosine, respectively; in CC_mut1_L, adenine at -9 bp from the dyad was replaced by cytosine; and in CC_mut1_R, cytosine at +10 bp from the dyad was replaced by guanosine. SHL: superhelix location. **(B)** Effect of temperature on the mobility of nucleosomal bands in native polyacrylamide gel electrophoresis. Nucleosomes migrated at low and high mobilities (denoted LM and HM, respectively). **(C)** Thermal stability assay of nucleosomes reconstituted from the 601 and CC_mut1 DNAs. A schematic diagram of nucleosome dissociation in a thermal stability assay is shown at the top of the thermal denaturation curves. The thermal denaturation curves of the nucleosomes reconstituted from the 601 or CC_mut1 DNA are red and blue, respectively. The native polyacrylamide gel electrophoresis of aliquots of the 601 and CC_mut1-nucleosomes incubated at 25 and 64°C is shown beside the thermal denaturation curves.

First, we analyzed the temperature-dependent changes in nucleosome position by native PAGE (**Fig. 4B**). As the temperature increased, nucleosomes were detected as three bands, shifting the band from low mobility (denoted LM in **Fig. 4B and C and Supplemental Fig. S10**) to high mobility (denoted HM in **Fig. 4B and C and Supplemental Fig. S10**). This result indicates that the position of the 601 nucleosome changes with increasing temperature, in good agreement with previous results (Sato et al. 2020). Compared to the control 601-nucleosome, the CC_mut1-nucleosome was clearly detected as three bands at 42°C, and the band with the highest mobility (HM) was predominant at and above 49°C. We also investigated the CC_mut1L- and CC_mut1R-nucleosomes to clarify the effects of individual mutations. The CC_mut1L- and CC_mut1R-nucleosomes both shifted to bands with higher mobility at lower temperatures than the control 601-nucleosome, consistent with the CC_mut1-nucleosome results. Thus, changing either one of the bases from A to C at position -9 or from C to G at position +10 in the 601-nucleosome caused the nucleosome position to become labile.

We next analyzed the thermal stabilities of the nucleosomes containing the 601 and CC_mut1 sequences. In this assay, the free histones dissociated from the nucleosome are detected by the binding of the fluorescent probe SYPRO Orange, and the nucleosomes are denatured by a biphasic process with increasing temperature: the first and second phases correspond to the H2A-H2B and H3-H4 dissociations, respectively (Taguchi et al. 2014; Sato et al. 2020) (**Fig. 4C**). The thermal stability profile of the CC_mut1 nucleosome overlapped perfectly with that of the 601-nucleosome. During the thermal denaturation process, aliquots of the 25°C and 64°C samples were analyzed by PAGE, showing that the positioning of the CC_mut1-nucleosome was more labile than that of the 601-nucleosome.

We further analyzed the major position of the nucleosome in the 193 bp fragment containing either the 601 or CC_mut1 sequences **(Supplemental Fig. S10**). First, the DNA bands with lower mobility for 601 and CC_mut1 nucleosomes (designated 601-LM and CC_mut1-LM, respectively) and those with higher mobility for the CC_mut1 nucleosomes (CC_mut1-HM) were purified by preparative PAGE (**Supplemental Fig. S10A**). After the nucleosomes were treated with MNase, the DNA fragments were purified and separated by native PAGE (**Supplemental Fig. S10B**). These fragments were further digested with the *Alu*I restriction endonuclease, and analyzed by native PAGE (**Supplemental Fig. S10C**). The DNA regions covered with histone proteins were determined by the migration properties of the *Alu*I-digested fragments. Based on these results, the primary location of the 601-LM and CC_mut1-LM nucleosomes was determined to include the central 601 sequence in a 193 bp fragment. In contrast, the position of the CC_mut1-HM nucleosome was found at the right end of the 193 bp fragment (**Supplemental Fig. S10D**). This implies that the CC_mut1 mutation caused a position shift to form a more translationally stable nucleosome, without affecting the thermal stability of the nucleosome itself.

## Discussion

Herein, we established a high-resolution mapping method for nucleosomes in the yeast genome, by analyzing a 139 bp fragment produced by chemical cleavage at positions ±70 *via* histone H2A-A122C. The major cleavage sites are consistent with the recent finding by Armeev et al. (Armeev et al. 2021) that the C-terminal tail of histone H2A inserts into the minor groove at the DNA ends in the nucleosome. Therefore, this method is unique in that it defines the location of the fully wrapped nucleosome in the yeast genome, since the 139 bp DNA fragments must have arisen from a nucleosome in the wrapped state.

It is noteworthy that comparisons of the distributions of dinucleotides within representative nucleosomes determined by H2A-A122C, H3-Q85C, and H4-S47C revealed a characteristic sequence bias at each chemical cleavage site. These biases may be due to the sequence preference of the copper chelating reagent inserted into the DNA minor groove and/or the local binding affinity of the histone to which the reagent is bound. The positional concordance of the representative nucleosomes determined by the three methods was not high, and the nucleosomes were distributed at intervals of 10 bp from the central peak of the positional concordance. This implies that the nucleosome is mobile among discrete translational positions spaced by 10 bp, which is consistent with prior studies (Meersseman et al. 1992; Buttinelli et al. 1993; Shen et al. 2001; Kim et al. 2006). Furthermore, hundreds of thousands of redundant nucleosomes were detectable in addition to the representative nucleosomes. Since the cleavage strength should correlate with the probability of nucleosome presence at a given location, the conformations of nucleosomes in the cell are considered to be highly dynamic. These dynamic features may be relevant to the recent report (Tan et al. 2023) that heterogeneous non-canonical nucleosomes, including alternative histone H2A-H2B conformations and partially dissociated DNA, predominate *in situ*.

Sequence-dependent nucleosome formation in the genome has been extensively analyzed in several organisms *in vivo* and *in vitro*, using nucleosomes reconstituted with natural and synthetic DNA fragments (Balasubramanian et al. 2009; Satchwell et al. 1986; Satchwell and Travers 1989; Albert et al. 2007; Kaplan et al. 2009; Travers et al. 2010; Cui and Zhurkin 2010; Brogaard et al. 2012a; Chereji et al. 2018; Lowary and Widom 1998). A general view of nucleosome sequence-dependence is that WW dinucleotides prefer the DNA minor-groove sites facing toward histone octamers with about 10 bp periodicity, while SS dinucleotides prefer the minor groove facing outward toward histones, as shown in chickens (Satchwell et al. 1986; Satchwell and Travers 1989), yeast (Albert et al. 2007; Ioshikhes et al. 2011; Brogaard et al. 2012a; Chereji et al. 2018), fruit flies (Mavrich et al. 2008), nematodes (Wright and Cui 2019), and human (Gaffney et al. 2012). Our H2A-A122C analysis is in good agreement with these previous reports and establishes the sequence-dependent formation into fully wrapped nucleosomes at high resolution *in vivo*.

Furthermore, we discovered the absence of consecutive G•C base-pairs ±9-11 bp from the dyad within the SHL ±1.0 major groove blocks; instead, the sites ±8-10 bp from the dyad favor WW dinucleotides. This tendency to avoid GG/CC base-pairs is also found in fission yeast and mice. Moreover, we showed that the introduction of GG/CC dinucleotides ±9-11 bp from the dyad in 601-reconstituted nucleosomes altered their positions in the 193 bp DNA fragment *in vitro*. To investigate the mechanistic reasons for this sequence dependence, we examined the atomic structures of the nucleosomes around SHL ±1.0 in previously solved crystal structures (Vasudevan et al. 2010; Frouws et al. 2016; Richmond and Davey 2003). In the 601 nucleosome (**Fig. 5**), the bases in the proximal strand (red) to histones at -9 and +9 are T and G, respectively. The side chain of arginine-40 (R40) penetrates into the minor groove, and its density is clearly detectable at -9, but not at +9. From a similar point of view, the bases in the proximal strand (red) at -9 and +9 are G and T, respectively, in the MMTV nucleosome (**Supplemental Fig. S11A**), while they are both A in the alpha-satellite nucleosomes (**Supplemental Fig. S11B**). In these three nucleosome structures, the densities of the H3-R40 side chain are detectable in the case of the A•T base pair at the ±9 bp positions, but less so for the G•C base pair in which G is in the proximal strand. Thus, a guanine residue in the proximal DNA strand could destabilize the H3-R40 insertion, while an A•T or T•A bp allows the stable insertion. This interpretation is consistent with recent MS simulations by Armeev et al.(Armeev et al. 2021), who found that the unwrapping of the nucleosomal DNA ends was coupled to the destabilization of the DNA near the dyad, through their mutual interactions with the portion of histone H3 (called H3-latch) between the H39 and R49 residues. Particularly, the loss of the H3-latch interactions with the DNAs was accompanied by the removal of H3-R40 from the DNA minor groove near the dyad. Thus, GG dinucleotides in the proximal strand at ±9-11 bp from the dyad may destabilize the interactions with H3-R40.

**Fig. 5.**
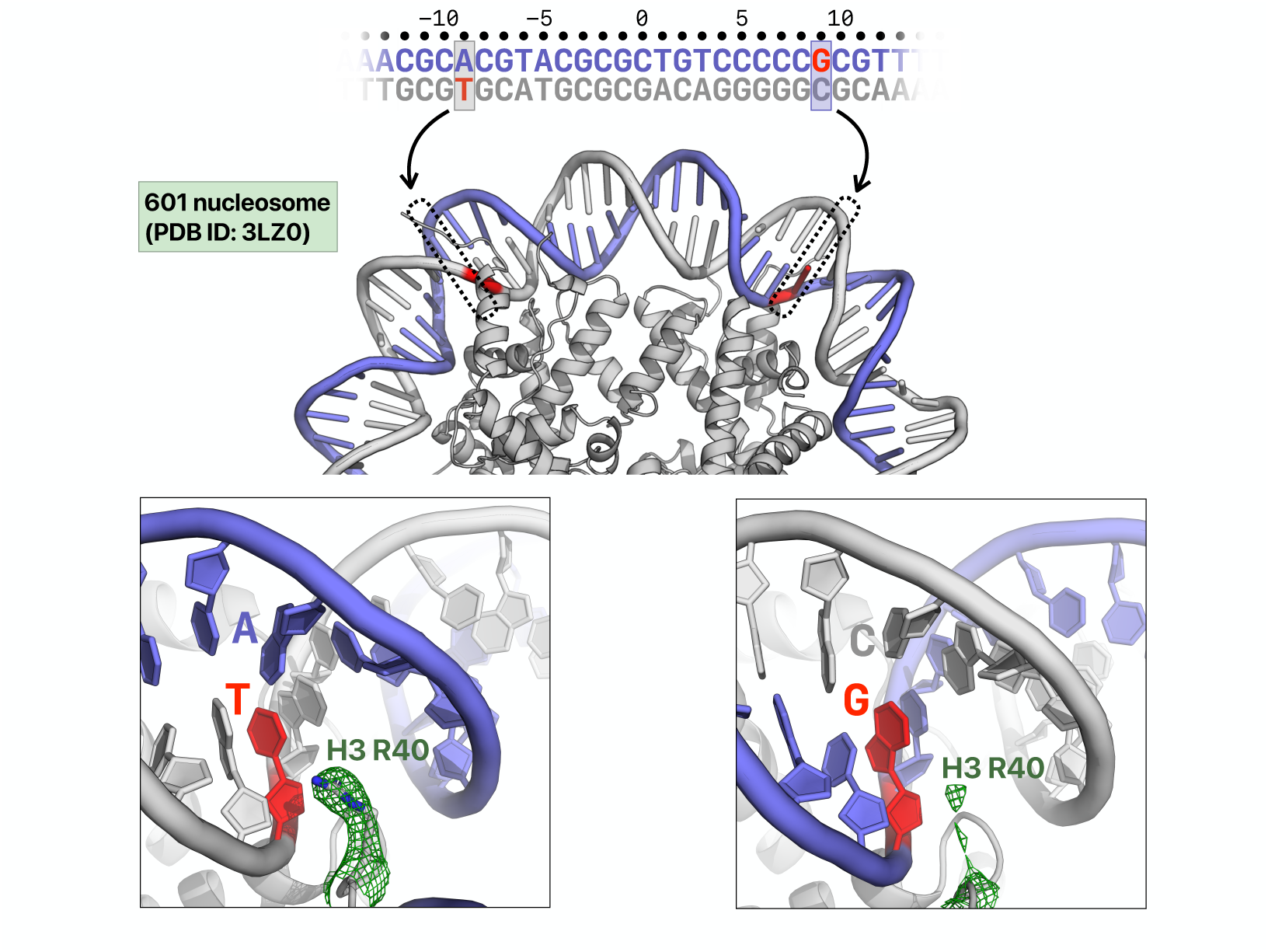
Crystal structure of the nucleosome composed of the 601 sequence around the dyad from the SHL -2 to +2 region. (PDB: 3LZ0, Vasudevan et al. 2010). The base pairs at -9 and +9 from the dyad as zero are highlighted by dashed boxes and gray shadowing in the top figures. Bases at the ±9 positions in the DNA strand proximal to the histone octamer are shown in red. The side chains of the histone H3 R40 residues are inserted into the minor grooves near ±9 bases, and the electron densities of the side chains of the R40 residues are shown in green shading. In all the structures, the electron density of the R40 side chain was clearly observed in the case of the A•T base pair at ±9, but it was ambiguous in the case of guanine in the proximal strand of DNA at ±9.

In summary, we established the sequence-dependency within nucleosomes at higher resolution in the yeast genome, and discovered a novel signature of nucleosomes: the exclusion of the GG dinucleotide at ±9-11 bp from the dyad in the proximal strand of the histone octamer. From this perspective, if spontaneous point mutations occur at these sites, then the position of the nucleosome could be significantly altered, with potentially significant consequences for gene expression and epigenetics. It also seems likely that encountering naturally occurring CC/GG-dinucleotides at positions ±9-11 bp in nucleosomal DNA would alter the fiber topology and wrapping state of the nucleosome, affecting the 3D genome organization as well as transcription factor binding, as reported recently (O’Dwyer et al. 2025; Wen et al. 2025).

## Materials and Methods

### Yeast strains

Yeast strains used were FY24 (a derivative of S288C) derivatives, with the genetic markers *MATα ura3-52 trp1Δ63 leu2Δ1*. To establish the H2A-A122C strain, pRS306 containing the *hta1* gene (-400 to +649) with the A122 mutation was constructed. Using the plasmid and *hta2Δ* strain, in which the *HTA2* gene in FY24 was replaced by the *KanMX* fragment, the H2A-A122C strain was constructed by two-step gene replacement. The genotype of H4-S47C strain, MHS3006 was [*MATα ura3-52 trp1Δ63 leu2Δ1 hhf1::S47C hhf2::KanMX*] (Fuse et al. 2017) (Supplemental Table S1).

### Isolation of yeast nuclei

Yeast nuclei were isolated as reported (Szent-Gyorgyi and Isenberg 1983; Shimizu et al. 1991). Yeast cells were cultured in 1L of SC medium until the OD_600_ reached 1.0–1.5, and NaN_3_ and phenylmethanesulfonyl fluoride (PMSF) were then added to the medium, at final concentrations of 0.13% and 0.5 mM, respectively. The cells were harvested, washed and treated with Zymolyase (10 mg/mL) in S-buffer (1.4 M D-sorbitol, 40 mM HEPES (pH 7.5), 0.5 mM MgCl_2_, 0.13% NaN_3_, 1 mM PMSF) for conversion to spheroplasts. The spheroplasts were homogenized in F-buffer [18% Ficoll™ PM400 (Cytiva), 20 mM PIPES (pH 6.5), 0.5 mM MgCl_2_, 1 mM PMSF] and the homogenate was layered onto 20 mL of GF-buffer [20% glycerol, 7% Ficoll™ PM400, 20 mM PIPES (pH 6.5), 0.5 mM MgCl_2_, 1 mM PMSF] and centrifuged for 30 min at 20,000xg. The pellet was resuspended in Ficoll buffer and centrifuged for 15 min at 3,500xg, and the resulting supernatant was centrifuged for 25 min at 20,000xg. The nuclei were obtained as the pellet.

### Site-directed chemical mapping

The site-directed chemical cleavages were performed as reported previously(Katsumata et al. 2021; Fuse et al. 2019, 2017). The nuclei pellet was resuspended in 4 mL of labeling buffer (1 M sorbitol, 50 mM NaCl, 10 mM Tris-HCl, pH 7.5, 5 mM MgCl_2_, 0.5 mM spermidine, 0.15 mM spermine). For labeling, 0.7 mM N-(1,10 phenanthroline-5-yl) iodoacetamide (Biotium cat# 92015) dissolved in dimethylsulfoxide was added to the nuclei suspension, to a final concentration of 0.14 mM of the labeling reagent. The labeling reaction was incubated with rotation at 4°C for 1 hour in the dark. After the reaction, the nuclei were recovered by centrifugation at 4,000xg for 10 min, and resuspended in 3 mL of labeling buffer. The nuclei were washed with labeling buffer four times, and resuspended in 3 mL of mapping buffer (1 M sorbitol, 2.5 mM NaCl, 50 mM Tris-HCl, pH 7.5, 5 mM MgCl_2_, 0.5 mM spermidine, 0.15 mM spermine). CuCl_2_ was added to the nuclei suspension to a final concentration of 0.01 mM, and the suspension was incubated for 5 min at room temperature. After centrifugation at 4,000xg for 10 min, the nuclei were washed three times with 3 mL of mapping buffer, and resuspended in 990 μL of mapping buffer. 3-Mercaptopropionic acid was added to a final concentration of 5.9 mM, and the suspension was incubated for 5 min at room temperature. The chemical reaction was initiated at room temperature by the addition of 0.4 M hydrogen peroxide to a final concentration of 6 mM, and at 10 and 20 min, aliquots of the reaction mixture (330 μL) were transferred to a new tube containing 2 μL of 0.5 M neocuproine to quench the reactions. The nuclei were pelleted and resuspended in 200 μL of mapping buffer, and treated with RNase and subsequently with proteinase K. The DNA was purified by phenol/CHCl_3_ extraction, followed by ethanol precipitation.

### MNase digestion of chromatin in isolated nuclei

The nuclei pellet was resuspended in 2 mL of digestion buffer [10 mM HEPES (pH 7.5), 0.5 mM MgCl_2_, 0.05 mM CaCl_2_, 1 mM PMSF]. The suspended nuclei were divided into 200 μL portions and digested at 37°C for 10 min, using successive 2-fold serial dilutions (64 to 8 units/mL) of MNase (Worthington), as reported (Shimizu et al. 1991). The DNA was purified in the same manner as in the chemical mapping described above.

### Parallel sequencing analysis

The chemically-cleaved (reaction for 20 min) or MNase-digested (64 units/mL) DNA samples described above were separated by 8% native polyacrylamide gel electrophoresis, and DNA fragments ranging from 50 to 250 bp in length were excised from the gel and purified. The ends of the chemically- or MNase-cleaved DNA were blunted using NEB’s Klenow Polymerase large fragment (cat# M0210L), followed by Lucigen’s DNA Terminator End Repair Kit (cat# 40035-2) (Brogaard et al. 2012b, 2012a). The paired-end library was prepared from the DNA fragments with a ThruPLEX® DNA-Seq Kit (Clontech, #R400674), and sequencing was performed using a NovaSeq/HiSeq sequencer.

### Datasets from previous studies

The unique and representative H4-S47C-based nucleosome maps liftovered for the sacCer3 coordinates were used (Fuse et al. 2017; Brogaard et al. 2012a). The total number of representative nucleosomes (67,510) was determined by subtracting 38 of the dyads at the chromosome ends from the number of original dyads (67,548). Raw reads for H3-Q85C-based fragments(Chereji et al. 2018) were obtained from SRA (http://www.ncbi.nlm.nih.gov/sra, SRP102823.

### Read mapping and processing of mapped data

Quality filtering and trimming (-t 5) of 50 mer reads were performed with fastp (0.19.7)(Chen et al. 2018). The 45 mer reads were mapped to the sacCer3 reference genome (R64-1-1, www.ensemble.org) with bwa mem (https://github.com/lh3/bwa). Properly paired reads were selected, split by chromosome names, and further split by mapped fragment length (from 20 to 280 bp) with sambamba (v0.6.6), viewed with the “proper_pair” and “template_length” options (https://lomereiter.github.io/sambamba/). For dyad identification, the pileup count of the 5’ end nucleotide was obtained with bedtools genomecov (https://bedtools.readthedocs.io/) with the options (-d -5 -strand + or -d -3 -strand -).

### Dyad calling, occupancy calculation and selection of representative nucleosomes

For the H4-S47C samples, the R package NuCMap (version 1.0, https://github.com/jipingw/NuCMap) was used for calling the dyads of unique and redundant nucleosomes. Properly mapped reads from two biological replicates were combined before dyad calling.

For the H2A-A122C and H3-Q85C samples, mapped fragments with 139±5 and 51±5 bp lengths were used for dyad calling. The number of fragments was counted and consecutively assigned to the genomic coordinates corresponding to the midpoint of the fragments. The sum of fragment numbers at each genomic coordinate was defined as the dyad score of the nucleosome located at that position.

The nucleosome occupancy was calculated by extending the dyad scores by 73 bp along the genome in both directions. Dyad score and occupancy data were stored as wiggle format files, binarized with igvtools (2.3.40), and visualized on the Integrative Genomics Viewer (IGV) browser (https://igv.org).

For the H2A-A122C and H3-Q85C samples, representative nucleosomes were selected as those with the highest dyad scores within the corresponding ±106 bp regions. Thus, the shortest distance between two neighboring representative nucleosomes should be 107 bp. Dyad-to-dyad distance was calculated as the distance between the dyads of representative nucleosomes.

### Measurement of positional redundancy (fuzziness) of nucleosomes

For a given representative nucleosome with a dyad base located at chromosomal coordinate *i*, all detected nucleosomes located within the surrounding 147-bp region (*i*-73 to *i*+73) were picked up. Then, the standard deviation for the relative positions of these nucleosomes relative to the representative dyad position (*i*) was calculated as a measure of the positional redundancy.

### Oligonucleotide frequency analysis

For the DNA sequences of the 201-bp regions centered at each representative nucleosome, reverse complement sequences were bundled to the forward strand sequences to symmetrize the frequency data. For grouping nucleosomes with respect to the dyad score, nucleosomes with the highest 1% and lowest 10% dyad scores were omitted, and the rest were split into nine groups (#1-9) in order of dyad score. The frequency of oligonucleotides starting from each nucleosomal coordinate was normalized by dividing the raw values by the genomic frequency. WW dinucleotide frequency was computed as the sum of normalized frequencies for “AA,” “AT,” “TA,” and “TT,” while for SS dinucleotide frequency, “CC,” “CG,” “GC,” and “GG” were used. frequencies for “AA,” “AT,” “TA,” and “TT,” while for SS dinucleotide frequency, “CC,” “CG,” “GC,” and “GG” were used.

### Reconstitution of nucleosomes *in vitro*

The 193 base-pair DNA fragment (601, CC_mut1, CC_mut1_L or CC_mut1_R) and human histone octamer were mixed in buffer, containing 10 mM Tris-HCl (pH 7.5), 2 M KCl, 1 mM dithiothreitol and 1 mM EDTA. Nucleosomes were reconstituted by the salt-dialysis method, in buffer containing 10 mM Tris-HCl (pH 7.5), 0.25 M KCl, 1 mM dithiothreitol, and 1 mM EDTA (Kujirai et al. 2018). The reconstituted nucleosomes were purified by a polyacrylamide gel electrophoresis-based method using a Prep Cell apparatus (Bio-Rad). To analyze the nucleosome positioning of CC_mut1_HM, the CC_mut1 nucleosome was heated at 55°C for 2 hours before purification. The purified nucleosomes were stored in buffer containing 20 mM Tris-HCl (pH 7.5) and 1 mM DTT.

### Nucleosome positioning analysis *in vitro*

Nucleosomes (35 ng/μL of DNA) containing 193 base-pair DNAs were heated in buffer, containing 10 mM Tris-HCl (pH 7.5) and 1 mM DTT, at the temperatures described in the Fig. 4B for 2 hours. Nucleosome repositioning was analyzed by nondenaturing 6% polyacrylamide gel electrophoresis in 0.2x TBE (17.8 mM Tris, 0.4 mM EDTA and 17.8 mM boric acid). After staining with ethidium bromide, the gel images were obtained using an Amersham imager (GE Healthcare). To determine the nucleosome positioning of 601_LM, CC_mut1_LM, and CC_mut1_HM, a restriction enzyme digestion assay coupled with MNase treatment was performed. The linker DNAs of 0.35 μg nucleosomes were digested with MNase (TAKARA, 0.8 units) at 37°C for 8 min in a 20 μL reaction solution, containing 20 mM Tris-HCl (pH 8.0), 10 mM Tris-HCl (pH 7.5), 25 mM NaCl, 2.5 mM CaCl_2_ and 1 mM dithiothreitol. The reactions were stopped by adding 10 μL of stop solution, containing 20 mM Tris-HCl (pH 8.0), 0.4 mg/mL Proteinase K (Roche), 0.1% SDS, 575 mM NaCl and 20 mM EDTA. The DNA fragments were separated by polyacrylamide gel electrophoresis and extracted with the Wizard SV gel and PCR clean-up system (Promega). The extracted fragments were digested by *Alu*I (New England Biolabs) and fractionated by polyacrylamide gel electrophoresis. The gel was stained with SYBR Gold, and the DNA fragments were detected by a Typhoon imager (GE Healthcare). The migration profiles were analyzed by the Image Gauge software (GE Healthcare), and the fragment lengths were estimated by 20 base-pair DNA markers as references.

### Thermal stability assay

The thermal stability of the nucleosomes containing 601 or CC_mut1 was assessed according to the previous report (Taguchi et al. 2014). The nucleosome (85 ng of DNA) was incubated in a solution containing 5x SYPRO Orange (Sigma-Aldrich), 20 mM Tris-HCl (pH 7.5) and 1 mM dithiothreitol. The SYPRO Orange fluorescence emitted by its binding to denatured histones was detected with a StepOnePlus Real-Time PCR system (Applied Biosystems). The temperature gradient ranged from 25°C to 95°C, in steps of 1°C/min. The fluorescence intensity was normalized relative to the fluorescence signal at 95°C.

### Data access

All raw sequencing data generated in this study have been deposited in the DDBJ Sequence Read Archive (DRA) under accession number PRJDB20610. The in-house scripts for this study are available at (https://github.com/hkatomed/KatoFuse2025).

## Acknowledgements

This work was supported in part by JSPS KAKENHI Grant numbers JP24K09416 to M.S., JP24K08832, JP23K27350 and JP21H00255 to H.Ka., JP16H06279(PAGS) to M.S. and H. Ka., JP24H02323, JP23H00372, JP22H04676 and JP22K19275 to Y.O., JP23H05475, JP23K24997 and JP23K28231 to W.K., JP24H02328 and JP23H05475 to H.Ku., JP25K09593 and JP22K06179 to S.S., JST ERATO grant JPMJER1901 to H.Ku., JST CREST grant JPMJCR24T3 to H.Ku., AMED BINDS JP22ama121017j0001 to Y.O., and JP25ama121009 to H.Ku. This work was also supported in part by the MEXT Promotion of Development of a Joint Usage / Research System Project: Pan-Omics DDRIC, MRCI for High Depth Omics, CURE: JPMXP1323015486 for MIB and RIIT in Kyushu University, and by Priority Research Funding from Meisei University.

## Author contributions

M.S., T.F., Y.K., and W.K. designed and established the H2A-A122C chemical mapping method, T.F. prepared chemical and MNase mapping samples for the NGS library, Y.O. prepared the NGS library and performed sequencing, H.Ka. and T.U. developed algorithms and performed bioinformatic analyses, S.S., H.Ku., M.S., and H.Ka. designed *in vitro* reconstitution experiments, S.S. and H.Ku. analyzed the reconstituted nucleosomes, W.K. analyzed the crystal structures of nucleosomes, and M.S. and H.Ka conceived, designed and supervised the work. M.S. and H.Ka. wrote the manuscript and all authors contributed to manuscript writing and editing.

